# Single-cell RNA sequencing reveals aortic cellular heterogeneity in MSC_ITGB3-treated ApoE-/- mice on a high-fat diet

**DOI:** 10.1101/2023.01.18.524536

**Authors:** Haijuan Hu, Demin Liu, Xue Geng, Mei Han, Wei Cui

**Author notes:** Corresponding author: (WC).

## Abstract

The aorta contains various cell types that are involved in the development of vascular inflammation and atherosclerosis. However, the cellular atlas of heterogeneous aorta cells, cellular responses, and intercellular communication has not been investigated in the background of a high-fat diet (HFD) and treatment with integrin beta 3-modified mesenchymal stem cells (MSC_ITGB3). In this study, 33,782 individual cells from mouse aortas under HFD with or without MSC_ITGB3 treatment were subjected to single-cell RNA sequencing as an unbiased analysis strategy. We generated a compendium of 30 different clusters, mainly smooth muscle cells (SMCs), endothelial cells, and immune cells. The proportion of the different cell types was considerably influenced by HFD and MSC_ITGB3. In the HFD group, genes associated with proliferation, migration, and collagen were highly expressed in the major SMC subpopulations. However, the expression of contraction-related genes in SMC subpopulations was significantly higher in the MSC_ITGB3 group than in the HFD group. After HFD consumption, subpopulations of ECs with active PI3K-Akt signaling pathway, ECM-receptor interaction, and contraction-related genes were significantly enriched. In the MSC_ITGB3 group, the number of dendritic cells (DCs), which are positively correlated with atherosclerotic lesion progression and contribute to lipid accumulation, and levels of inflammatory factors notably decreased. Our findings provide data on the composition, signaling pathways, and cellular communication of the aorta cells following stem cell treatment as well as on the evolution and progression of atherosclerotic disease. The findings may help in improving the treatment of atherosclerosis.

## Introduction

Atherosclerosis is a characteristic of vascular inflammation, and its complications are one of the leading causes of mortality worldwide [1]. Atherogenesis involves the interaction between local and global pro- and anti-inflammatory factors. In recent years, great progress has been made in the treatment of atherosclerosis, including systemic pharmacological treatment and percutaneous coronary intervention [2, 3]. However, the incidence of atherosclerotic complications, such as stroke and myocardial infarction, remains relatively high [4].

Mesenchymal stem cell (MSC) transplantation brings new light to atherosclerosis treatment [5-7]. Targeted modification drives MSCs to their destination and improves repair at the site of injury. Overexpression of fibroblast growth factor 21 significantly increases the migration and homing of MSCs to injured brain tissues [8]. C-X-C chemokine receptor type 5 (CXCR5) modification enhanced the migration ability of MSCs towards CXCL13 in a mouse model of contact hypersensitivity, leading to decreased levels of inflammatory cell infiltration and proinflammatory cytokine production [9]. The integrin family of receptors enables cells to interact with their microenvironment [10, 11]. The main recognition system for cell adhesion is constituted by integrin receptors, along with proteins containing Arg-Gly-Asp (RGD) attachment sites [12, 13]. The primary sequence of integrin beta 3 (ITGB3), a highly conserved region in all beta subunits of integrin, is referred to as the RGD-cross-linking region [14]. Consequently, we genetically engineered stem cells to overexpress ITGB3 via *in vitro* lentiviral transduction. However, whether stem cell migration to the plaque site affects the vascular cellular composition and heterogeneity *in vivo* remains unclear. To date, few studies have investigated the changes in the composition and heterogeneity of vascular cells in atherosclerotic plaques following MSC therapy.

In this study, we aimed to elucidate the effects of stem cell therapy on atherosclerotic vascular cell composition. Accordingly, we used single-cell RNA sequencing (scRNA-seq) to investigate cell heterogeneity and differential functional states within the aortic wall of mice fed a high-fat diet (HFD) and treated with or without ITGB3-modified MSCs (MSC_ITGB3) (MSC_ITGB3 and HFD groups, respectively). Collectively, we demonstrated that a high number of MSC_ITGB3 can migrate to the injury site and promote plaque repair following injection into the mouse tail vein. Furthermore, we performed differential analyses of lineage heterogeneity and functional changes and elucidated the underlying vascular cell communication mechanisms from the aortic wall in the HFD and MSC_ITGB3 groups. Cluster analysis results revealed 30 clusters and 5 distinct cell types. Importantly, compared with HFD aortas, aortas treated with MSC_ITGB3 exhibited changes in the cell subsets, transcriptome characteristics, and biological functions.

## Materials and methods

### Data collection and processing

Single-cell data and samples with data type *Mus musculus* were divided into HFD and MSC_ITGB3 groups (n=3 per group). All experiments were performed on ApoE-/-mice fed an HFD. In the MSC_ITGB3 group, MSC_ITGB3 cells were injected four times via the tail vein (1 × 10^6^ cells/injection every week), starting at week 9, and aortic sampling was performed at the end of week 12 for scRNA-seq. A single-cell suspension of aortic cells was prepared using a previously described enzymatic digestion protocol [15].

The atherosclerosis Bulk-seq dataset GSE43292 [1] was downloaded from the Gene Expression Omnibus database (https://www.ncbi.nlm.nih.gov/geo/). It comprised *Homo sapiens* samples, and its assay platform was GPL6244. The dataset was derived from 32 patients with hypertension. Each patient provided one sample of an atherosclerotic plaque containing the core of the shoulder of the plaque (Stary classification type IV and above) and one sample of a distant macroscopically intact tissue (type I or II), resulting in a total of 64 samples.

### Quality control of atherosclerosis data using Seurat

R (version 4.1) and the ‘Seurat’ R package (version 4.0.5) [16] were used. Seurat objects for each sample were created from single-cell data and then merged using the “merge” function. The proportion of mitochondrial genes to all inherited genes is a major factor that determines whether a cell is in a steady state. A cell with a higher proportion of mitochondrial genes than all other genes is generally considered to be in a stress state.

Therefore, we removed cells with > 20% mitochondrial genes. As low-quality cells or empty droplets usually have few genes and double cells may exhibit an abnormally high number of genes, we also removed cells with features < 200 or > 7,000. Ultimately, 33,782 cells were obtained (Fig 1).

**Fig 1.**
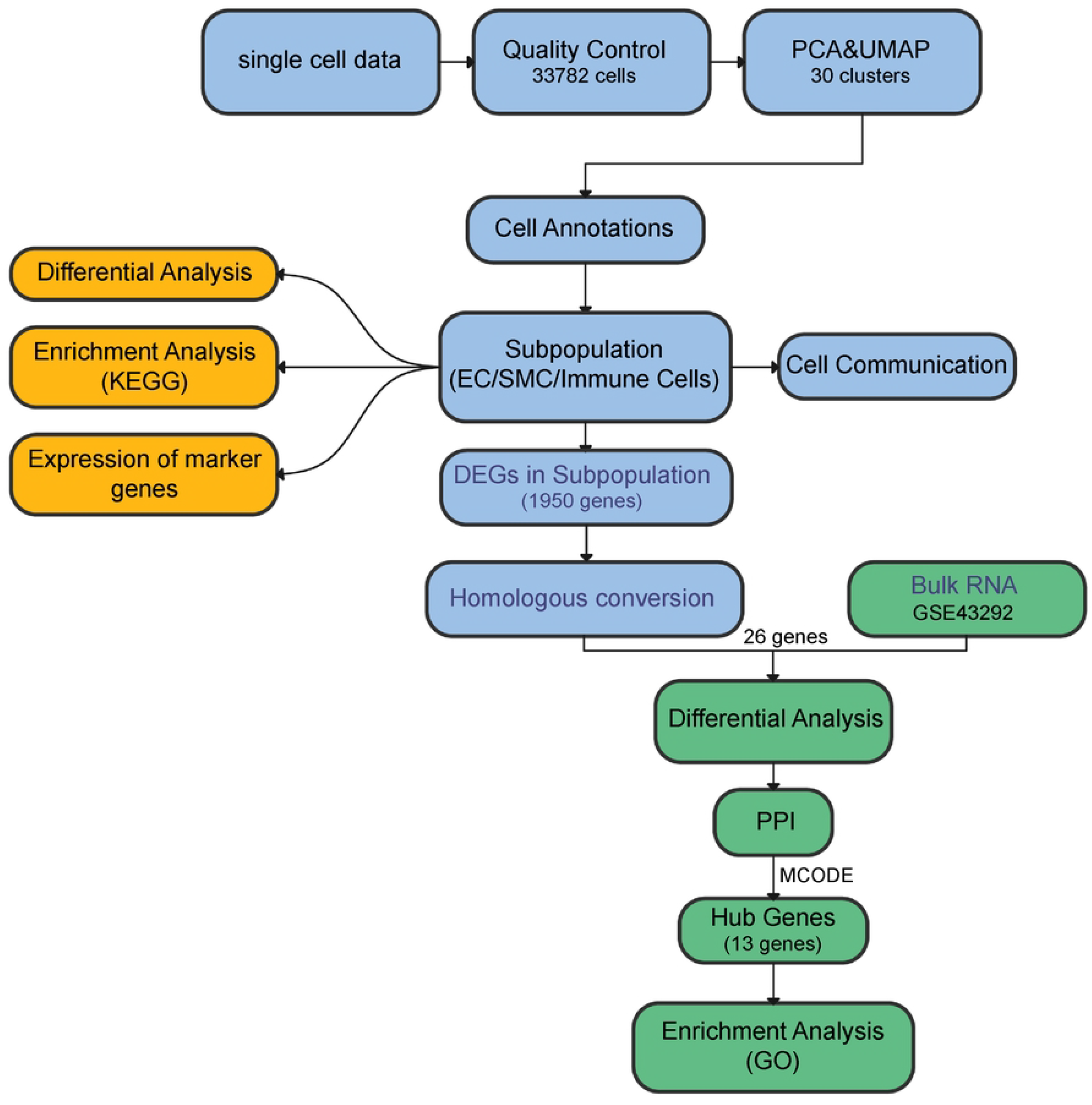
Project flowchart. We developed a flowchart of bioinformatic analysis to regularize and characterize different data.

The sequencing depth of the dataset was normalized using the “NormalizeData” function with the default “LogNormalize” normalization method. Subsequently, 2,000 variable features of the dataset were detected by calling the “FindVariableFeatures” function using the “vst” method. The data were then scaled using “ScaleData” to exclude the effects of sequencing depth. The “Elbowplot” function was used for the Principal component analysis (PCA) [17], to identify the significant principal components (PCs) and visualize the *p*-value distribution. Subsequently, the data were de-batched using the “RunHarmony” function. Finally, 24 PCs were selected for uniform manifold approximation and projection (UMAP) analysis. The default parameters of “FindNeighbors” and 22 PC dimension parameters were used to construct the Euclidean distance-based K-nearest neighbors in the PCA space. The Louvain algorithm was used to optimize the class groups by calling the “FindClusters” function, which divides the cells into 30 different clusters with a resolution of 0.8. Finally, the “RunUMAP” function was used for dimensionality reduction to visualize and investigate the dataset.

### Gene enrichment analysis

Gene Ontology (GO) [18] enrichment analysis is a common approach for large-scale functional enrichment studies of genes in different dimensions and at different levels. It is generally performed at three levels: molecular function, biological process, and cellular component. Kyoto Encyclopedia of Genes and Genomes (KEGG) [19] is a widely used database for storing data on genomes, biological pathways, diseases, and drugs. The “clusterProfiler” (version 4.2.0) [20] R package was used to perform GO and KEGG functional annotation of differentially expressed genes between the macroscopically intact tissue and atheroma plaque, in the bulk RNA data, and between cells in single-cell data, to assess significantly enriched biological processes. The significance threshold was set at *p* < 0.05, and the results were visualized using bar graphs.

### Cell annotation

Cell types were identified based on their marker genes [21]: *Pecam1* and *Cdh5* for endothelial cells (ECs), *C1qb* and *Lyz2* for immune cells, *Col1a1* and *Col1a2* for fibroblasts, *Tagln* and *Myh11* for smooth muscle cells (SMCs), and *Kcan1* and *Plp1* for neural cells.

### SMC subpopulation annotation

The method described above was used for the dimensionality reduction and clustering of SMCs. Twelve principal components were determined as statistically significant inputs to UMAP and were divided into 12 clusters. Based on marker gene expression [21], SMCs were classified into three cell subpopulations: SMC_1 (*Fn1, Ctgf*, and *Eln*); SMC_2 (*Myl6, Acta2*, and *Tagln*); and SMC_3 (*Gpx3, Colec11*, and *Col6a1*). The default Wilcox test of the FindAllMarkers function was then used to determine differentially expressed genes between different cell types (logfc.threshold = 0.25).

### EC subpopulation annotation

The same method described above was used for dimensionality reduction and EC clustering. Twelve principal components were determined as statistically significant inputs to UMAP and were divided into 11 clusters. Based on marker gene expression [21], ECs were classified into three cell subpopulations: EC_1 (*Cytl1, Cpe, Clu*, and *Pam*); EC_2 (*Fabp4, Ly6c1, Sparcl1*, and *Igfbp7*); and EC_3 (*Mmrn1, Fgl2, Igfbp5*, and *Lbp*). The default Wilcox test of the FindAllMarkers function was then used to determine differentially expressed genes between different cell types (logfc.threshold = 0.25).

### Immune cell subpopulation annotation

The same method described above was used for the dimensionality reduction and clustering of immune cells. Sixteen principal components were selected as statistically significant inputs to UMAP and were divided into 15 clusters. Based on marker gene expression [21], immune cells were classified into three cell subpopulations: monocytes/macrophages (*Cd68, C1qb*, and *Lyz2*); dendritic cells (DCs; *H2-Ab1* and *H2-Eb1*); and T cells (*Cd3d, Cd3g*, and *Nkg7*). The default Wilcox test of the FindAllMarkers function was then used to determine differentially expressed genes between different cell types (logfc.threshold = 0.25).

### Cell communication analysis

The ‘CellChat’ (http://www.cellchat.org/) [22] R package was used to analyze cell-cell communication networks from scRNA-seq data. Single-cell expression profiles were combined with known ligands, receptors, and their cofactors to calculate the strength of interactions in cell-cell communication. Network analysis and pattern recognition methods were used to predict the major incoming and outgoing signals in the cells as well as the coordination between these cells and signals.

### Key gene screening from bulk RNA data

The “homologene” (version 1.4.68.19.3.27) R package was used for the homologous transformation of single-cell data for differential genes between SMCs, ECs, and immune cells. Subsequently, the expression of these homologous genes in the 64 samples was determined from the GSE43292 dataset using the “limma” (version 3.50.0) [23] R package to perform differential analysis on the resulting new macroscopically intact tissue and atheroma plaque groups, from which differential genes were extracted (*p* < 0.05, |logFC| > 1).

The STRING online database (https://string-db.org/) [24] was used to analyze the interactions between differentially expressed genes. Protein–protein interaction (PPI) networks were constructed using Cytoscape [25] (version 3.9.1) to map the interactions between the functions of the proteins, including direct physical interactions and indirect functional correlations. The cytoHubba [26] plugin was used to assign values to each gene using a topological network algorithm to sort and discover key genes and sub-networks.

## Results

### Single-cell data-based cell type annotation reveals a high degree of cellular heterogeneity in samples

A total of 33,782 cells were obtained after filtering based on quality control criteria and normalization of the scRNA-seq data (Fig 2A). Overall, 2,000 highly variable genes were selected for subsequent analysis, and the top 10 genes were annotated (Fig 2B). PCA was performed to identify usable dimensions and screen for relevant genes, and 22 PCs were selected for subsequent analysis. Using the UMAP dimensionality reduction, the cells were divided into 30 separate clusters (Fig 2C, D). Thirty clusters were identified using marker genes for each cell type. Clusters 2, 3, 5, 7, 8, 9, 10, 11, 15, 21, 22, 24, 25, 26, 27, 28, and 29 had a total of 16,717 cells annotated as Fibroblast, accounting for 49.48% of all cells.Clusters 1, 12, 13, 17, 18, and 20 had 6,402 cells annotated as Immune_cell, accounting for 18.95% of all cells. Clusters 0, 14, and 23 had 5,090 cells annotated as Endothelial_cell, accounting for 15.07% of all cells. Clusters 4, 6, and 16 had a total of 5,090 cells annotated as SMCs, accounting for 15.07% of all cells. Cluster 19 had 483 cells and was annotated as Neural, accounting for 1.43% of all cells (Fig 2E).

**Fig 2.**
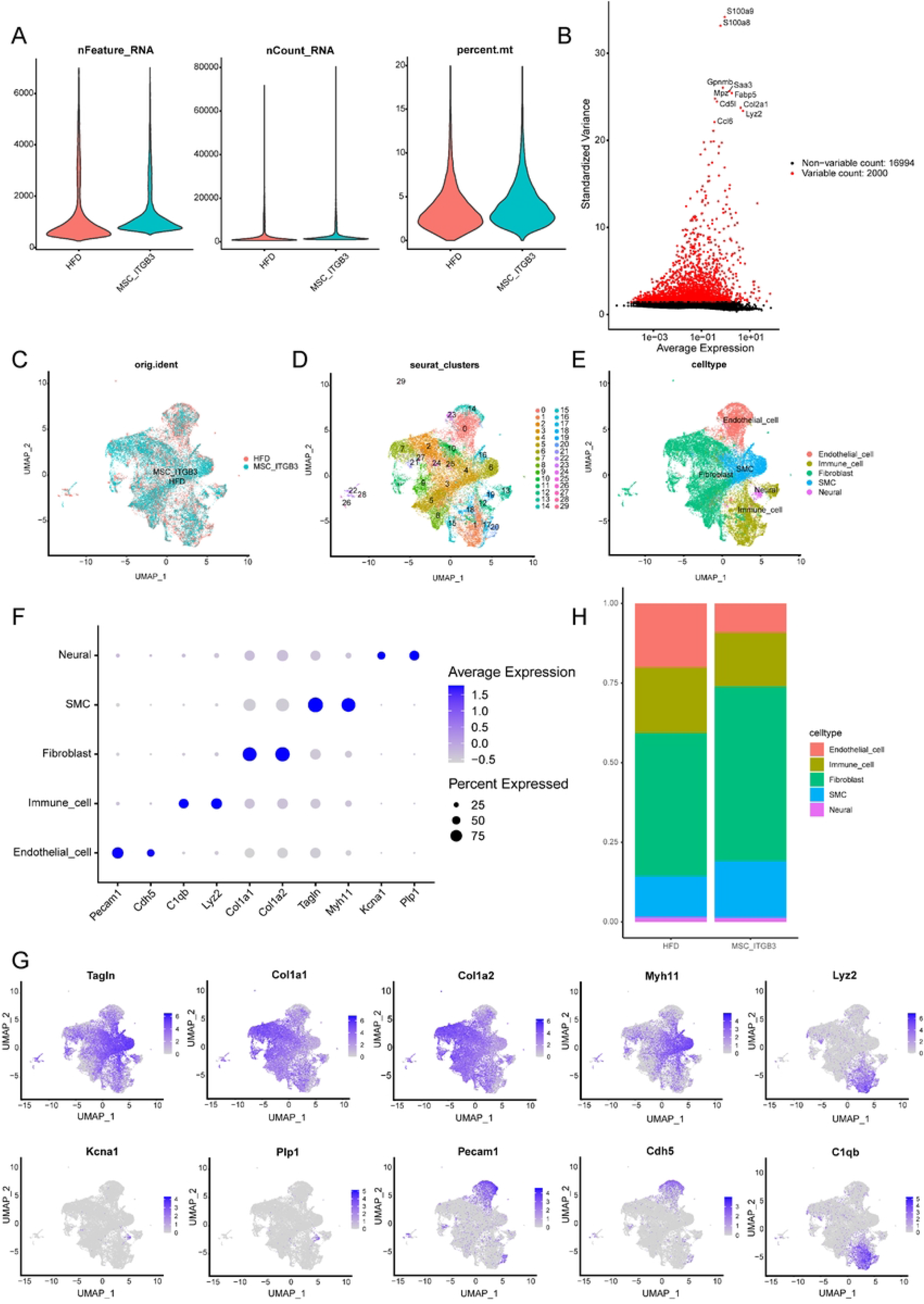
Quality control and characterization of the single-cell dataset. (A) Two samples from the single-cell dataset were selected, and 33,782 cells were included in the analysis following quality control. (B) Standard deviation scatter plot demonstrating high gene variability in cells, with the top 10 genes labeled. Cluster analysis was performed on single-cell data samples and colored based on sample (C), cluster (D), and cell type (E). (F) Prominent marker genes for each cell type (Endothelial_cell, Immune cell, Fibroblast, SMC, and Neural). (G) Cell type proportions in the single-cell datasets of the high-fat diet (HFD) and MSC_ITGB3 groups. (H) Expression profiles of the prominent marker genes for each cell type.

The specific marker genes of each cell type were used to construct a dot plot (Fig 2F) and feature map (Fig 2G), and the percentage of each cell type in the HFD and MSC_ITGB3 groups was calculated (Fig 2H).

### SMC subpopulation analysis

Further cluster analysis of SMCs revealed three subpopulations (Fig 3A). In the HFD group, SMC_2 accounted for 74.17% of the SMCs, and SMC_1 and SMC_3 together accounted for 25.83%. In the MSC_ITGB3 group, SMC_2 accounted for 79.07%, and SMC_1 and SMC_3 together accounted for 20.93% (Fig 3B). Differential analysis (*p* < 0.05) of these three SMC subpopulations was then performed, and the top 10 differential genes of each cell subpopulation were used to construct a heat map of differential gene expression (Fig 3C). Finally, KEGG enrichment analysis was performed for each SMC subpopulation, and enrichment of the following pathways was found: focal adhesion, diabetic cardiomyopathy, prion disease, proteoglycans in cancer, and Alzheimer’s disease in SMC_1 (Fig 3D); focal adhesion, diabetic cardiomyopathy, prion disease, proteoglycans in cancer, and chemical carcinogenesis (reactive oxygen species) in SMC_2 (Fig 3E); and extracellular matrix (ECM)-receptor interaction, focal adhesion, human papillomavirus infection, phosphoinositide 3-kinase (PI3K)-protein kinase B (Akt) signaling pathway, and hypertrophic cardiomyopathy in SMC_3 (Fig 3F).

**Fig 3.**
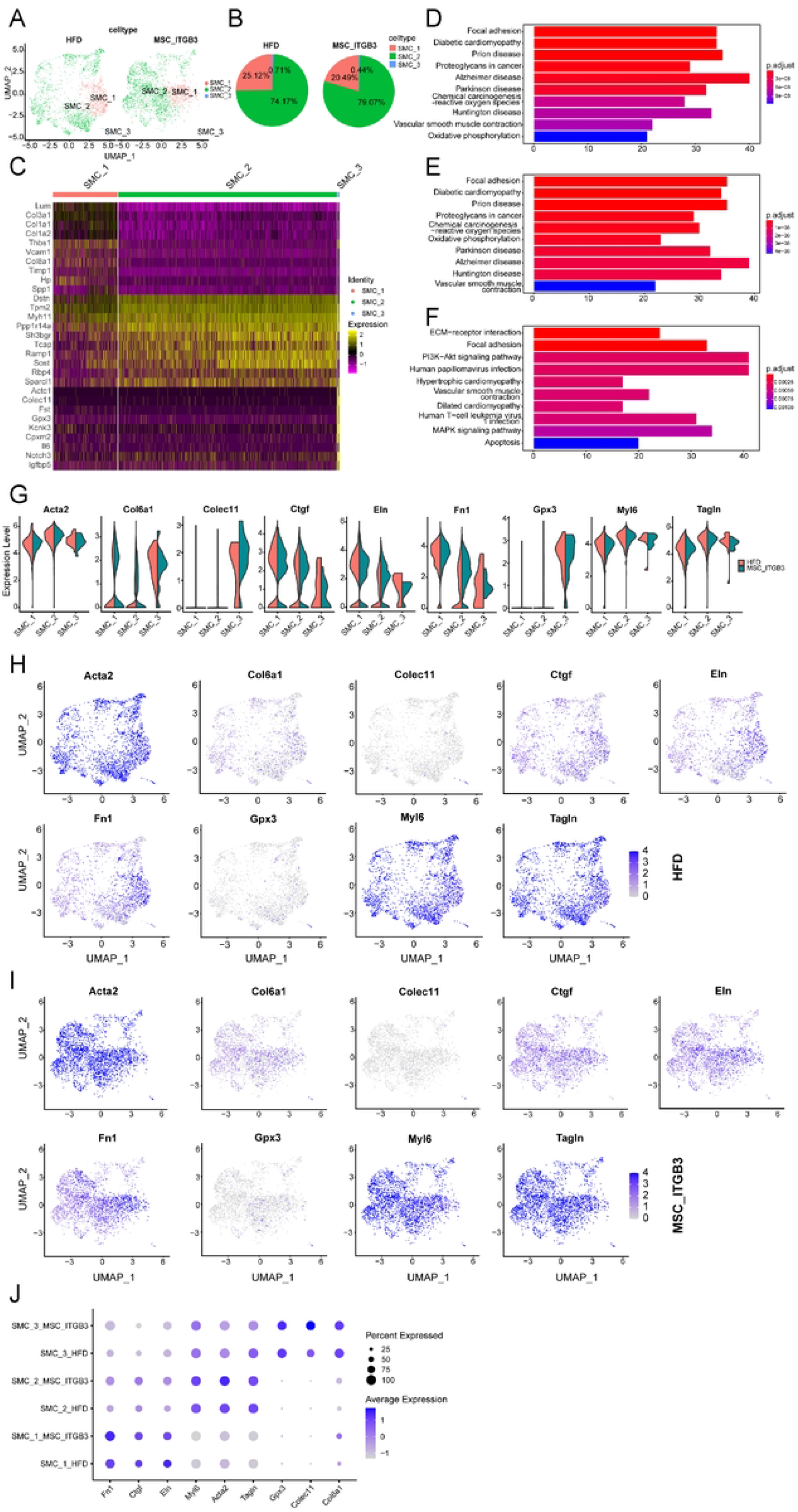
Comparison of SMC subpopulations in the HFD and MSC_ITGB3 groups. (A) UMAP plots of SMC subpopulations (SMC_1, SMC_2, and SMC_3) from the HFD (2,400 cells) and MSC_ITGB3 (2,733 cells) groups. (B) Percentage of SMC subpopulations in the HFD (SMC_1, 25.12%; SMC_2, 74.17%; SMC_3, 0.71%) and MSC_ITGB3 (SMC_1, 20.49%; SMC_2, 79.07%; SMC_3, 0.44%) groups. (C) Heat map of the top 10 marker genes for each subpopulation. (D–F) Top 10 KEGG-enriched pathways obtained using differentially expressed genes from the SMC_1 (D), SMC_2 (E), and SMC_3 (F) subpopulations. (G) Violin plots of selected marker gene expression in SMC subpopulations in both HFD and MSC_ITGB3 samples. (H) Feature map of the expression of selected marker genes of the SMC subpopulations in the HFD group. (I) Feature map of the expression of selected marker genes of the SMC subpopulations in the MSC_ITGB3 group. (J) Expression of specific genes in the SMC subpopulations.

Marker genes specific to each SMC subpopulation were selected (*Fn1, Ctgf, Eln, Myl6, Acta2, Tagln, Gpx3, Colec11*, and *Col6a1*) and grouped together to construct violin plots (Fig 3G), feature maps (Fig 3H and 3I), and dot plots (Fig 3J) to represent subpopulation-specific gene expression in the HFD and MSC_ITGB3 groups. High expression of the following genes was observed: genes involved in proliferation and migration (*Fn1, Ctgf, Eln*) in SMC_1, genes related to contractile markers (*Myl6, Acta2, Tagln*) in SMC_2, and collagen and redox genes (*Gpx3, Colec11, Col6a1*) in SMC_3.

### EC subpopulation analysis

Further cluster analysis of the ECs revealed three subpopulations (Fig 4A). In the HFD group, EC_1 accounted for the largest proportion (92.93 %), followed by EC_2 (5.79%) and EC_3 (1.28%), respectively. In the MSC_ITGB3 group, EC_1 accounted for 96.53%, followed by EC_2 (1.06%) and EC_3 (2.41%), respectively (Fig 4B). Differential analysis (*p* < 0.05) of these three EC subpopulations was then performed, and the top 10 differential genes of each subpopulation were used to construct a heat map of differential gene expression (Fig 4C). Finally, KEGG enrichment analysis for each EC subpopulation was performed, and enrichment of the following pathways was found: fluid shear stress and atherosclerosis, focal adhesion, proteoglycans in cancer, PI3K-Akt signaling pathway, and ECM-receptor interaction in EC_1 (Fig 4D); ECM-receptor interaction, coronavirus disease 2019 (COVID-19), fluid shear stress and atherosclerosis, focal adhesion, and proteoglycans in cancer in EC_2 (Fig 4E); and focal adhesion, AGE-RAGE pathway in diabetic complications, fluid shear stress and atherosclerosis, proteoglycans in cancer, and mitogen-activated protein kinase signaling pathway in EC_3 (Fig 4F).

**Fig 4.**
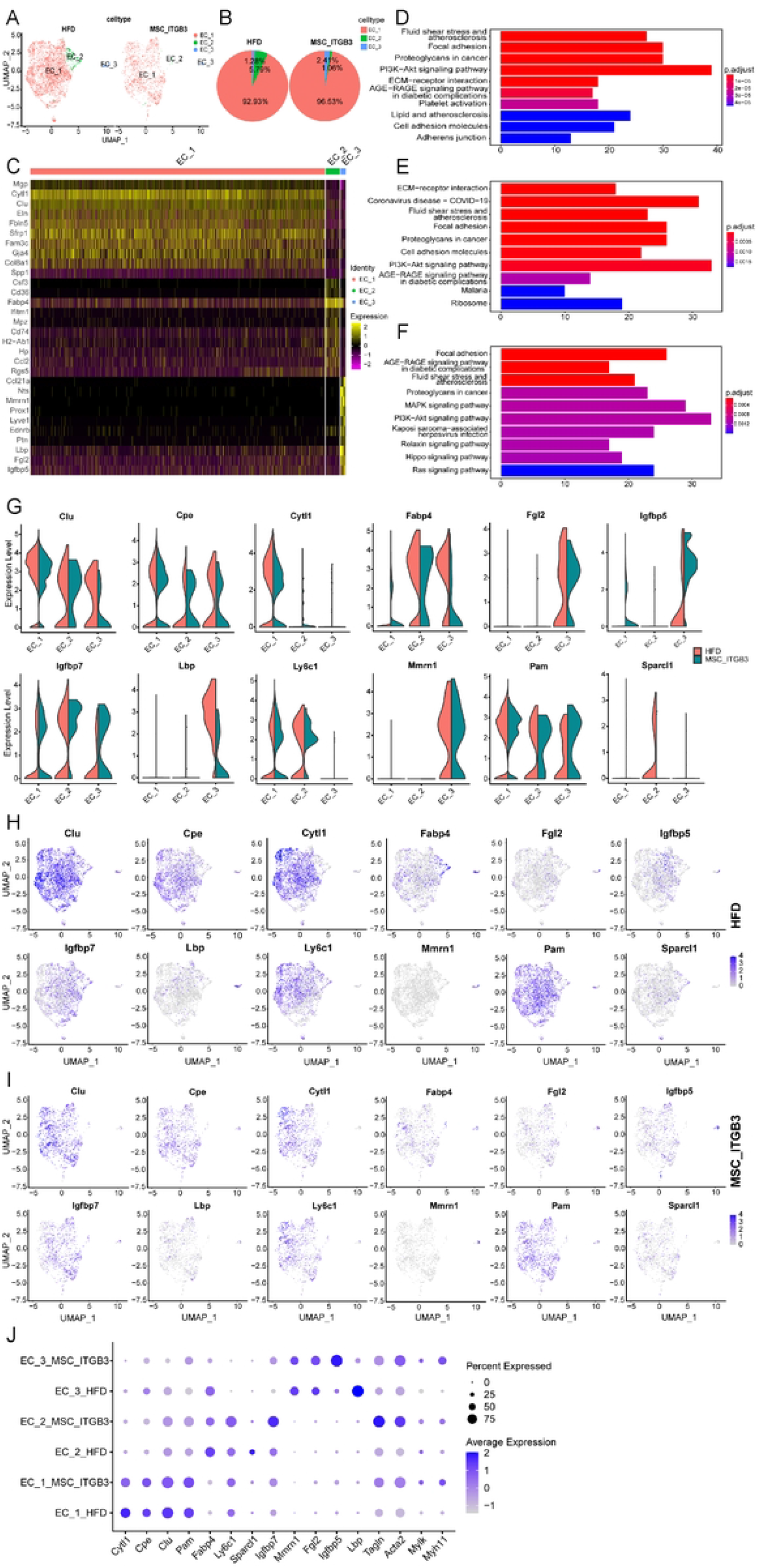
Comparison of EC subpopulations in the HFD and MSC_ITGB3 groups. (A) UMAP plots of EC subpopulations (EC_1, EC_2, and EC_3) from the HFD (3679 cells) and MSC_ITGB3 (1411 cells) groups. (B) Percentage of EC subpopulations in the HFD (EC_1, 92.93%; EC_2, 5.79%; EC_3, 1.28%) and MSC_ITGB3 (EC_1, 96.53%; EC_2, 1.06%; EC_3, 2.41%) groups. (C) Heat map of the top 10 marker genes for each subpopulation. (D-F) Top 10 KEGG-enriched pathways obtained using differentially expressed genes from the EC_1 (D), EC_2 (E), and EC_3 (F) subpopulations. (G–I) Violin plots (G) and feature maps (H and I) constructed through the selection and grouping of marker genes (*Cytl1, Cpe, Clu, Pam, Fabp4, Ly6c1, Sparcl1, Igfbp7, Mmrn1, Fgl2, Igfbp5*, and *Lbp*) specific to each EC subpopulation. Specific marker genes as well as contractility-driving genes (*Tagln, Acta2, Mylk*, and *Myh11*) were used to represent the specific gene and contractility-associated gene expression in the three EC subpopulations in the HFD and MSC_ITGB3 groups (J).

Marker genes (*Cytl1, Cpe, Clu, Pam, Fabp4, Ly6c1, Sparcl1, Igfbp7, Mmrn1, Fgl2, Igfbp5*, and *Lbp*) specific to each EC subpopulation were selected and grouped together to construct violin plots (Fig 4G) and feature maps (Fig 4H and 4I). Specific marker genes as well as contractility genes (*Tagln, Acta2, Mylk, Myh11*) were used to represent the specific gene and contractility-associated expression in the three EC subpopulations in the HFD and MSC_ITGB3 samples (Fig 4J). Genes associated with contractility were expressed at significantly higher levels in the MSC_ITGB3 group than in the HFD group.

### Immune cell subpopulation analysis

Further cluster analysis of the immune cells revealed three subpopulations (monocytes/macrophages, DCs, and T cells; Fig 5A). In the HFD group, monocytes/macrophages accounted for the largest proportion of immune cells (69.14%), followed by DCs (18.95%) and T cells (11.91%), respectively. In the MSC_ITGB3 group, monocytes/macrophages accounted for the largest proportion of immune cells (69.40%), followed by T cells (17.56%) and DCs (13.05%), respectively (Fig 5B). Differential analysis (*p* < 0.05) of these immune cell subpopulations was then performed, and the top 10 differential genes of each subpopulation were used to construct a heat map of differential gene expression (Fig 5C). Finally, KEGG enrichment analysis for each immune cell subpopulation was performed, and enrichment of the following pathways was found: Lysosome, Rheumatoid arthritis, Tuberculosis, Phagosome, and Antigen processing and presentation in DCs (Fig 5D); COVID-19, Ribosome, Lysosome, Rheumatoid arthritis, and phagosome in monocytes/macrophages (Fig 5E); and COVID-19, Ribosome, phagosome, rheumatoid arthritis, and Th17 cell differentiation in T cells (Fig 5F).

**Fig 5.**
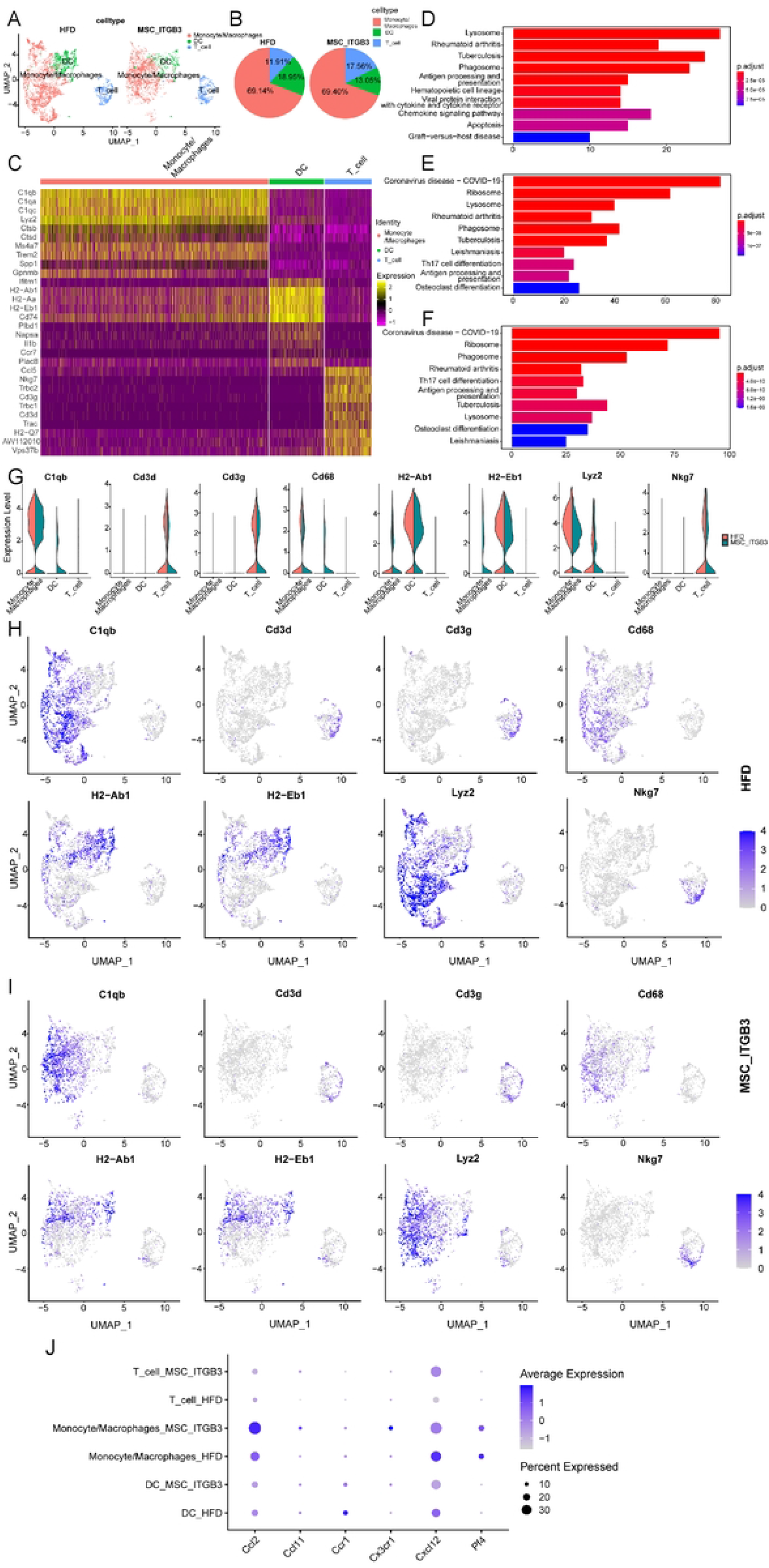
Comparison of immune cell subpopulations in the HFD and MSC_ITGB3 groups. **(A)** UMAP plots of immune cell subpopulations (monocytes or macrophages, DCs, and T cells) from the HFD (3,788 cells) and MSC_ITGB3 (2614 cells) groups. (B) Percentage of immune cell subpopulations in the HFD (monocyte and macrophages, 69.14%; DCs, 18.95%; T cells, 11.91%) and MSC_ITGB3 (monocyte and macrophages, 69.40%; DCs, 13.05%; T cells, 17.56%) groups. (C) Heat map of the top 10 marker genes for each subpopulation. (D-F) Top 10 KEGG-enriched pathways obtained using genes differentially expressed in the monocytes or macrophages (D), DC (E), and T cell (F) subpopulations. (G) Violin plots of selected marker gene expression in immune cell subpopulations in HFD and MSC_ITGB3 samples. (H) Feature map of expression of selected immune cell subpopulation marker genes in the HFD group. (I) Feature map of expression of selected immune cell subpopulation marker genes in the MSC_ITGB3 group. (J) Expression of inflammatory factor-encoding genes in the immune cell subpopulations.

Marker genes specific to each immune cell subpopulation were selected (*C1qb, Cd3d, Cd3g, Cd68, H2-Ab1, H2-Eb1, Lyz2*, and *Nkg7*) and grouped together to construct violin plots (Fig 5G) and feature maps (Fig 5H and 5I) to visualize subpopulation-specific gene expression. The expression of inflammatory factors (Ccl2, Ccl11, Ccr1, Cx3cr1, Cxcl12, and Pf4) in the immune cells of the HFD and MSC_ITGB3 groups are expressed as dot plots (Fig 5J). Among them, proinflammatory cytokines were highly expressed in monocytes/macrophages.

### Cell–cell communication

The total number of interactions between EC, SMC, and immune cell subpopulations (Fig 6A) as well as a complete interaction weight plot (Fig 6B) were obtained using ‘CellChat.’ Sankey diagrams were constructed to illustrate the cells interacting with each cell subtype during incoming (Fig 6C) and outgoing communication (Fig 6D) and the underlying signaling pathways. The diagrams show that compared with the other subpopulations, SMC_2, SMC_3, EC_1, EC_3, and monocytes/macrophages had a higher number of interactions with other cell types. Additionally, monocytes/macrophages had the strongest intensity of interaction with other cell types.

**Fig 6.**
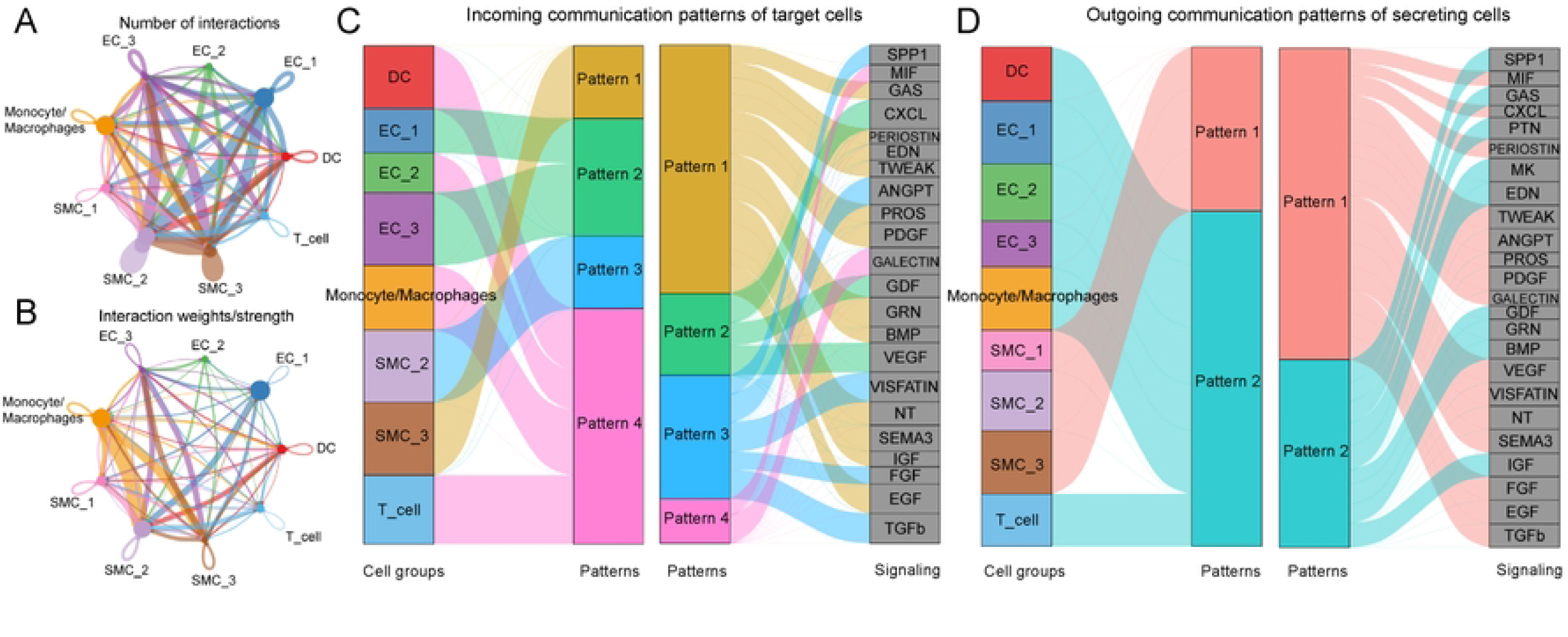
Ligand-receptor interaction analysis to assess communication between different cell subpopulations within atherosclerotic samples. (A) Circular plot depicting the communication network between cells in intracellular SMC, EC, and immune cell subpopulations (the number of cells in each cell type is proportional to the size of the circle; line widths indicate the number of signal interactions). (B) Circular plot demonstrating the intensity of interactions between cells contained in intracellular SMC, EC, and immune cell subpopulations (line widths indicate the intensity of cell-cell interactions). Sankey plots showing intercellular interactions during incoming (C) and outgoing (D) communication and the signaling pathways involved.

### Differential analysis of single-cell transcriptomic data

Differential analysis (*p* < 0.05) of ECs, SMCs, and immune cells was performed, and the top 10 differential genes in each subpopulation were used to generate a heat map (Fig 7A) of differential gene expression. The top five differential genes among the three cell subpopulations were used for feature map construction using UMAP (Fig 7B-7D).

**Fig 7.**
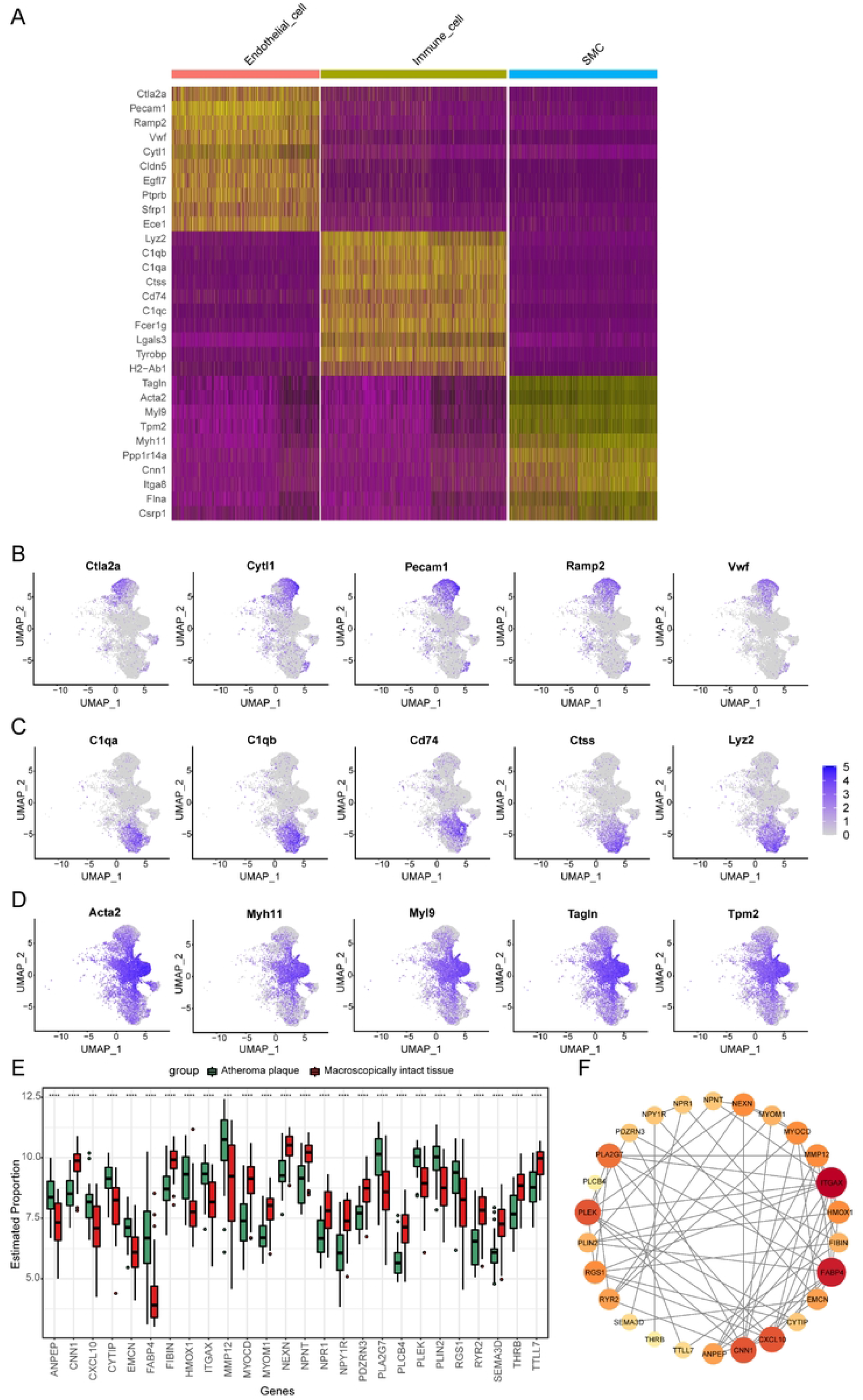
Analysis of differentially expressed genes in EC, SMC, and immune cells of HFD mice and MSC_ITGB3 mice. (A) Heat map of top 10 differentially expressed genes in EC, immune cell, and SMC subpopulations. (B–D) Feature maps constructed through selection of top five differentially expressed genes in EC (B), immune cell (C), and SMC (D) subpopulations. (E) Box plot of 26 differentially expressed genes from atheroma plaque and macroscopically intact tissues. (F) PPI network for differentially expressed genes. *p < 0.05; **p ≤ 0.01; ***p ≤ 0.001; ****p ≤ 0.0001.

Next, the differential genes of ECs, SMCs, and immune cells were transformed to their human homologs and screened in the GSE43292 transcriptome dataset for differential analysis. Twenty-six differential genes (*FABP4, MMP12, HMOX1, PLA2G7, PLIN2, ANPEP, PLEK, CYTIP, RGS1, ITGAX, CXCL10, EMCN, MYOM1, NEXN, TTLL7 THRB, NPNT, NPR1, PDZRN3, SEMA3D, CNN1, FIBIN, PLCB4, RYR2, NPY1R*, and *MYOCD*) were obtained. Box-line plots were plotted to illustrate the differential genes between atheroma plaque and macroscopically intact tissue samples (Fig 7E). The interactions between the differentially expressed genes were analyzed using STRING database and visualized using the Cytoscape software (Fig 7F).

Thirteen hub genes with the greatest interaction between the differentially expressed genes (*MMP12, CXCL10, HMOX1, ANPEP, ITGAX, FABP4, RGS1, PLEK, CNN1, RYR2, NEXN, MYOM1*, and *MYOCD*; Fig 8A) were extracted using the MCODE algorithm in the cytoHubba plugin. GO functional enrichment analysis of these 13 hub genes was performed, and negative regulation of vascular-associated SMC proliferation, muscle system processes, negative regulation of SMC proliferation, regulation of vascular-associated SMC proliferation, and vascular-associated SMC proliferation were found to be significantly enriched in these 13 hub genes (Fig 8B).

**Fig 8.**
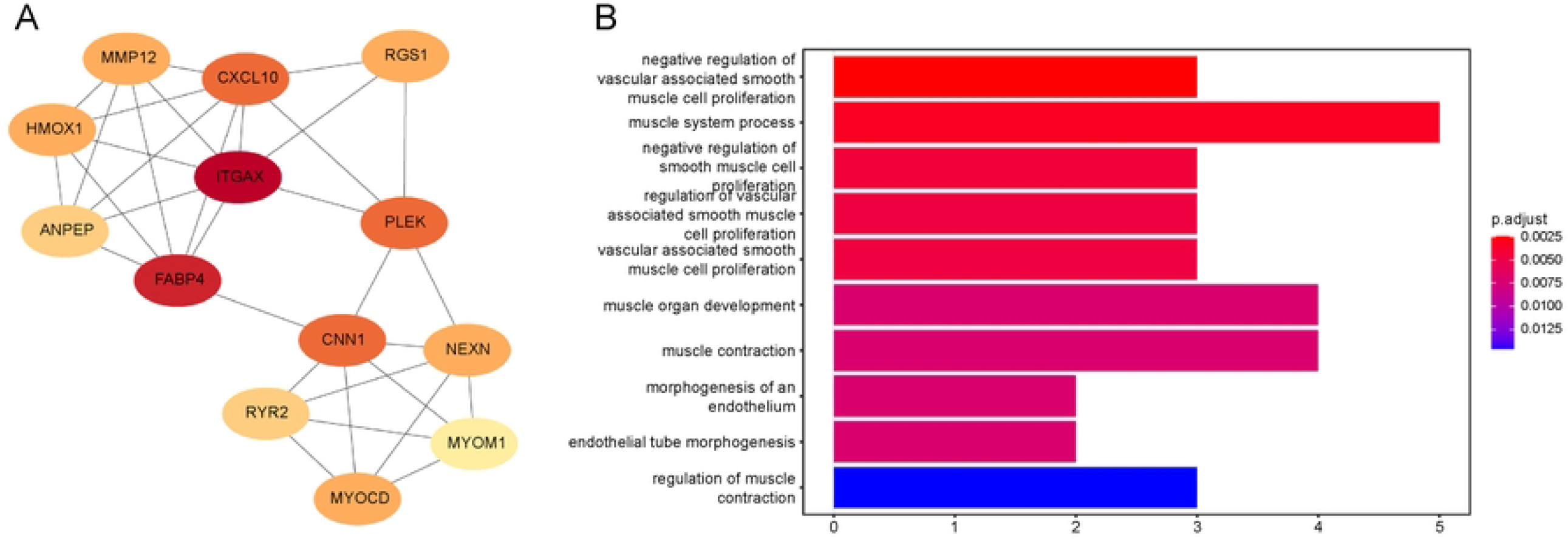
Hub gene screening and enrichment analysis. (A) Graph of hub gene interaction intensity calculated using the MCODE algorithm in the cytoHubba plugin. (B) Top 10 GO-enriched pathways obtained using 13 hub genes.

## Discussion

Owing to increasing incidence of risk factors such as hypertension, hyperlipidemia, and diabetes, atherosclerotic disease remains the leading cause of death. Stem cell therapy provides a new direction for atherosclerosis treatment. scRNA-seq technologies have been designed to reveal the heterogeneity of vascular cells, including those comprising normal vessels, those in ApoE^-/-^ mice, and those with aneurysms [21, 27, 28]. Many multifunctional cell populations related to cardiovascular diseases are composed of vascular tissue.

Heterogeneity in cell morphology and gene expression distinguishes between the main cell phenotypes and defines multiple subgroups with different functions. However, the heterogeneity and transcriptional features of the vascular cells of atherosclerotic aorta associated with stem cell therapy have not been explored. In this study, we used the latest scRNA-seq technology to comprehensively show the aortic cellular composition of - mice fed an HFD and provide new insights into the altered gene expression profiles in MSC_ITGB3-treated aortic cells.

We described the genes and signaling pathways expressed in 33,782 cells from the whole aorta and identified multiple subtypes in SMCs, ECs, and immune cells, suggesting that these cells include multiple functional populations in the aortic wall. Our data identified three subpopulations of SMCs in the aorta: synthetic (SMC_1), contractile (SMC_2), and inflammatory (SMC_3) SMCs. These subtypes are consistent with those identified by previous reports [27, 29]. The proliferating SMC cluster expresses several synthetic marker genes. The SMC_1 cluster expresses several proliferation and migration marker genes (*Fn1, Ctgf*, and *Eln*). The decrease in the synthetic gene subpopulation SMC_1 is an important condition that reverses the pathological progression of SMC phenotypic transformation in MSC_ITGB3-treated mice [30]. Cells in this cluster may play an adaptive role in vascular tissues with their migration function and high proliferation rate. SMC_2 highly expressed contractile transcription factors (*Myl6, Acta2*, and *Tagln*) under HFD conditions. Moreover, the expression of contraction-related genes in the MSC_ITGB3 group was significantly increased, indicating that the vasoconstrictive function of blood vessels improved after stem cell treatment. In addition, the inflammatory subgroup SMC_3 exhibited the highest expression of collagen and oxidation genes and the lowest expression of contractile genes. Stem cell therapy promoted the downregulation of proteinases and proinflammatory cytokines, including *Fn1, Ctgf, Eln Gpx3, Colec11, and Col6a1*. Thus, our sequencing data from HFD and MSC_ITGB3 mice demonstrates phenotypic diversity of vascular SMC and provides considerable insight into the heterogeneity of SMC in HFD- and MSC_ITGB3-treated vessels.

Our current study also provides a detailed analysis of EC gene expression signatures following stem cell therapy. ECs are known to have many subtypes. In our analysis, we identified three distinct EC profiles in the atherosclerotic aorta, and their gene expression profiles showed different functional characteristics. EC_1 accounts for the largest proportion of all EC subgroups and may play a key role in disease development. The PI3K-Akt signaling pathway and ECM-receptor interaction were significantly enriched in EC_1. PI3K-PKB/Akt is a highly conserved signaling pathway, and its activation mediates many cellular functions, including angiogenesis, survival, growth, transcription, proliferation, and apoptosis [31, 32]. ECMs interact with cell surface receptors to regulate cell behavior, such as cell communication, proliferation, migration, and adhesion [33, 34]. Specifically, cytokine-like protein 1 (CYTL1), a classical secretory protein, was downregulated in the MSC-ITGB3 group. CYTL1 is involved in neutrophil activation and the generation and release of reactive oxygen species during pathogenic infection [35]. Downregulation of *Cytl1* indicates a reduction in the inflammatory response following stem cell therapy. This difference indicates the functional characteristics of EC_1 in inflammation and adhesion. Based on the differential gene expression profiles, the EC_2 and EC_3 subgroups express genes that contribute to EC adhesion and leukocyte migration.

Atherosclerosis is a vascular inflammatory disease that involves the influx, proliferation, and activation of immune cells [36]. Several studies have demonstrated the heterogeneity of plaque cells and the proinflammatory effects of non-foam macrophages [37, 38]. However, the effect of stem cell therapy on the heterogeneity of immune cells in atherosclerotic plaques has not been reported. Based on our sequencing data, we identified three major subpopulations in the whole aorta: monocytes/macrophages, DCs, and T cells.

Among them, monocytes/macrophages, which accounted for the largest proportion of immune cells in the HFD and MSC_ITGB3 groups, play an important role in the phagosome pathway and COVID-19. Lysosome and antigen presentation signaling pathways were mainly enriched in DCs. The number of DCs is positively correlated with atherosclerotic lesion progression and contributes to lipid accumulation and disease initiation and progression [39]. Moreover, DCs can secrete a variety of proinflammatory factors, including tumor necrosis factor, interleukin (IL)-6, and IL-1β. After MSC_ITGB3 treatment, the number of DCs significantly decreased and expression levels of inflammatory factors considerably reduced. Hence, our data revealed characteristic changes in immune cells in the MSC_ITGB3-treated vascular aorta. scRNA-seq analysis provides a reliable tool for studying cell-to-cell interactions. By analyzing intercellular communication, we showed that the interactions of ECs, SMCs, and monocytes/macrophages trigger complex intercellular communication pathways that are directly or indirectly involved in the regulation of atherosclerosis. However, owing to limited funding, scRNA-seq was not performed on the unmodified MSC treatment group in this study, and the difference in treatment between unmodified MSCs and MSC_ITGB3 could not be compared.

In summary, our analysis comprehensively revealed the transcriptomic profile of atherosclerotic mouse aorta after MSC_ITGB3 treatment. After applying dimensionality reduction and clustering analysis, several functionally distinct candidate subpopulations were identified from the atherosclerotic vessels. These findings demonstrate the cellular diversity in plaques and provide insights into the cellular composition of the treated aorta and the function of individual cell subtypes.

## Acknowledgements

We would like to acknowledge Editage (https://www.editage.com/) for English language editing.

## Notes

### Competing Interest Statement

The authors have declared no competing interest.

